# Perinuclear force regulates SUN2 dynamics and distribution on the nuclear envelope for proper nuclear mechanotransduction

**DOI:** 10.1101/2022.12.16.520837

**Authors:** Jiahao Niu, Xiaotian Wang, Wenxue Zhao, Yao Wang, Yizhi Qin, Xiaoshuai Huang, Boxin Xue, Cheng Li, Yujie Sun

## Abstract

The trans-luminal LINC (Linker of Nucleoskeleton and Cytoskeleton) complex plays a central role in nuclear mechanotransduction by coupling the nucleus with cytoskeleton. High spatial density and active dynamics of LINC complex have hindered its precise characterization for the understanding of underlying mechanisms how the linkages sense and respond to mechanical stimuli. In this study, we focus on SUN2, a core component of LINC complex interconnecting the nuclear lamina and actin cytoskeleton and apply single molecule super-resolution imaging to reveal how SUN2 responds to actomyosin contractility. Using stochastic optical reconstruction microscopy (STORM), we quantitated the distribution pattern and density of SUN2 on the basal nuclear membrane. We found that SUN2 undergoes bidirectional translocation between ER and nuclear membrane in response to actomyosin contractility, suggesting that dynamic constrained force on SUN2 is required for its proper distribution. Furthermore, single molecule imaging unveils interesting dynamics of SUN2 molecules that are regulated by both actomyosin contractility and laminA/C network, whereas SUN2 oligomeric states are not affected by actomyosin contractility. Lastly, the mechanical response of SUN2 to actomyosin contractility was found to regulate expression of mechano-sensitive genes located in lamina-associated domains (LADs) and perinuclear heterochromatin. Taken together, our results reveal how SUN2 responds to mechanical cues at the single-molecule level, providing new insights into the mechanism of nuclear mechanotransduction.

## Main

Nuclear mechanotransduction defines the process in which the nucleus senses the mechanical stimuli derived from extracellular or cytoskeleton then transformed into intranuclear biochemical signals^1,2^. Distinct from the traditional biochemical signaling and directional cargo transportation, mechanical signaling can be transmitted more quickly through the cytoskeleton^3,4^. Over the past two decades, nuclear mechanotransduction has emerged as an important player in cell migration^5^, nuclear localization^6,7^, DNA damage response^8^, and chromatin organization^9,10^. Despite of the observations, the underlying mechanisms of nuclear mechanotransduction are poorly understood. So far, several mechanisms have been proposed for nuclear mechanotransduction: mechanical stimuli trigger permeability alterations of nuclear pore complexes (NPCs) and calcium fluxion ^11–13^; force triggers chromatin reorganization ^14,15^; force leads to changes of nuclear membrane protein post-translational modification levels^16^; force transduction through the perinuclear proteins^17^. As hundreds of proteins with diverse functions are localized on the nuclear envelope, how they function during nuclear mechanotransduction has become an urgent question to solve.

LINC complex is a key mechanotransduction element on the nuclear membrane, which consists of SUN (Sad1 and UNC-84) proteins on the inner nuclear membrane (INM) interacting with KASH (Klarsicht, ANC-1, and Syne/Nesprin homology) proteins on the outer nuclear membrane (ONM)^18–20^. In mammalian somatic cells, SUN1-containing LINC complex is associated with microtubule *via* the mediation of molecular motors. In contrast, SUN2-containing LINC complex is directly connected to the perinuclear actin through the calponin homology (CH) domain of Nesprin and assembles a linear array called TAN (transmembrane actin-associated nuclear) lines^6,21–23^. Inside the INM, SUN2 interacts with nuclear lamina through the N-terminal domain^24,25^, thereby physically connecting the actin cytoskeleton and the nucleoskeleton. SUN2 has been reported to play a role in cell migration and cancer inhibition^26,27^. Several previous studies have also revealed the roles of SUN2-containing LINC complex in nuclear mechanical response. For instance, Nesprin-1 and Nesprin-2 deficiency were found to cause nuclear envelope and lamin deformations^28^. Another study showed that SUN2 defects can block force-induced chromatin stretching^4^. A recent work characterized several mechano-sensitive phosphosites of SUN2 in the C terminal responding to cyclic tensile strain^29^. While these studies have established the essential role of SUN2-containing LINC complex as a load bearing structure, as a physical linkage with the cytoskeleton, lacking the characterization for SUN2 at the high spatiotemporal resolution has hindered the understanding of underlying mechanisms how the linkages sense and respond to mechanical stimuli.

In this study, we focused on how SUN2 responds to actomyosin contractility at the single molecule level and then regulates downstream gene expression. Given its high spatial density, we applied STORM super-resolution imaging to quantitate the distribution of SUN2 and found that the constraint force on SUN2 by actomyosin contractility is required for its proper distribution on the nuclear membrane and ER. Furthermore, single molecule tracking revealed that SUN2 dynamics were both regulated by actomyosin contractility and laminA/C. Interestingly, the oligomeric state of SUN2 does not change under different actomyosin contractility conditions. Finally, we found SUN2 can contribute to the structural organization of perinuclear chromatin and regulate expression of mechano-sensitive genes. Taken together, our work sheds light on the underlying mechanism by which the LINC component protein SUN2 responds to mechanical cues at the single molecule level, providing new insights into the understanding of nuclear mechanotransduction.

## Results

### 2.1 Actomyosin contractility level regulates the ratio of SUN2 distributed on the nuclear envelop and ER

The distribution pattern of nuclear envelope (NE) protein is an important prerequisite for its biological functions ^30–32^. To this end, we first investigated how actomyosin contractility affects the distribution of SUN2 on the nuclear membrane. We used RhoA activator to promote and Y27632 to suppress actomyosin contractility respectively, which hereinafter are referred to as RhoA group and Y27632 group for short. Given that the high spatial density of SUN2 on NE is beyond the resolving power of conventional microscopy, we applied STORM imaging, a super-resolution imaging mode that offers both high spatial resolution and single molecule counting capability, to image immunofluorescently labeled SUN2 on the flat basal side of the nuclear membrane. We observed that the density of SUN2 became higher in RhoA group and lower in Y27632 group in comparison with the wild-type cells, indicating that actomyosin contractility affects the distribution of SUN2 molecules on NE **(Fig. 1a)**. We defined the nearest neighbor distance (NND) between SUN2 clusters to quantify the sparsity of SUN2. The data show that the density of SUN2 in the RhoA group (NND = 0.86±0.10 μm) was significantly higher than that in the Y27632 group (NND=1.66 ±0.61 μm) **(Fig. 1b and Extended Data Fig. 1a)**. With the high spatial resolution of STORM imaging, we were able to resolve individual SUN2 clusters on NE. Importantly, both the size of SUN2 clusters and the expression level of SUN2 were found to remain similar under different actomyosin contractility levels **(Fig. 1c, Extended Data Fig. 1b, c)**. These data suggest that actomyosin contractility may promote recruitment of SUN2 molecules to NE without affecting the expression nor the oligomerization of SUN2. We then applied conventional wide field fluorescence imaging of SUN2 on endoplasmic reticulum (ER) as ER surrounds and connects with the nuclear membrane. SUN2 was found to be distributed on both ER and NE with similar density in the untreated cells. Interestingly, we found that SUN2 was almost completely relocated from ER to NE under high level of actomyosin contractility. In contrast, SUN2 was more enriched at ER when suppressing actomyosin contractility **(Fig. 1d, e)**. Taken together, these results indicated that SUN2 is distributed on ER and NE with a certain proportion under cellular homeostasis and dynamic actomyosin contractility regulates the ratio of SUN2 distributed on NE and ER.

**Fig. 1.**
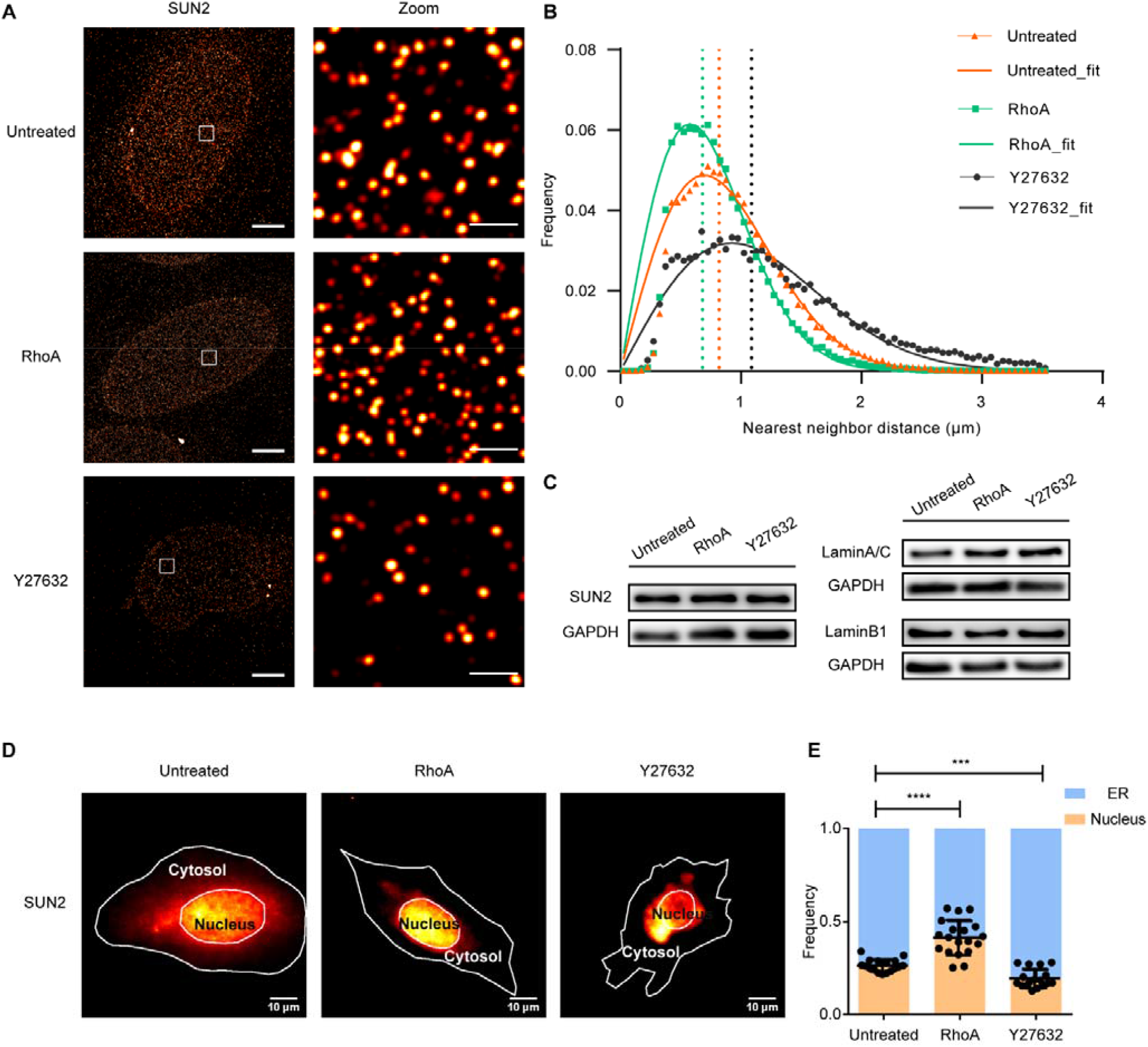
Actomyosin contractility regulates SUN2 distribution on the nuclear membrane. (**a**) Representative STORM imaging of SUN2 under untreated, RhoA, Y27632 conditions, respectively. Magnified views of the regions in the white boxes as right. Scale bar, 20 μm and 2 μm in the original and magnified images, respectively. (**b**) Histograms of the nearest neighbor distance for all clusters in untreated, RhoA, Y27632 conditions. The dashed line represents the median value, untreated: 0.8191 μm, RhoA: 0.6809 μm, Y27632: 1.0875 μm, bin size, 0.05. (**c**)Western blot of SUN2, LaminA/C, LaminB1 in whole cell extracts under RhoA and Y27632 treatments. (**d**) Representative fluorescence images of the distribution of SUN2 on the NE and cytosol (ER) under untreated (n=17 cells), RhoA (n=19 cells), Y27632 (n=17 cells) conditions. White lines represent cell and nucleus boundaries. Scale bar 10 μm. (**e**) Proportion of SUN2 between NE and cytosol (ER), ***, P < 0.0005; ****, P < 0.0001.

### 2.2 Actomyosin contractility regulates SUN2 dynamics on nuclear envelope

Nuclear mechanotransduction is a fairly dynamic process which converts the force from cytoskeleton into signals inside the nucleus. To further explore the role of SUN2 during this dynamic process, we applied Halotag-based ^33^ single-molecule tracking system in MB-MDA-231 cells to investigate the dynamic properties of SUN2 molecules under different actin mechanical conditions. Exogenous Halotag-fused SUN2 was expressed in SUN2 knockout cells to avoid the interference of various SUN2 isoforms **(Extended Data Fig. 2)**. Single molecule trajectories of individual SUN2 molecules showed that compared to the untreated group, the dynamics of SUN2 became more constrained in the RhoA group while more dispersed in the Y27632 group **(Fig. 2a and Supplementary Video 1-3)**. We performed mean squared displacement (MSD) analysis of the single molecule trajectories and found that SUN2 molecules clearly fell into two subgroups: a more mobile group and a more constrained group (18% constrained in untreated). In the RhoA group, the proportion of the constrained subgroup was increased compared to the mobile subgroup (45% constrained); In contrast, the proportion of the constrained subgroup was decreased compared to the mobile subgroup in the Y27632 group (9% constrained) **(Fig. 2b and Extended Data Fig. 3a-c)**. These data indicated that actomyosin contractility can regulates SUN2 dynamics on NE.

**Fig. 2.**
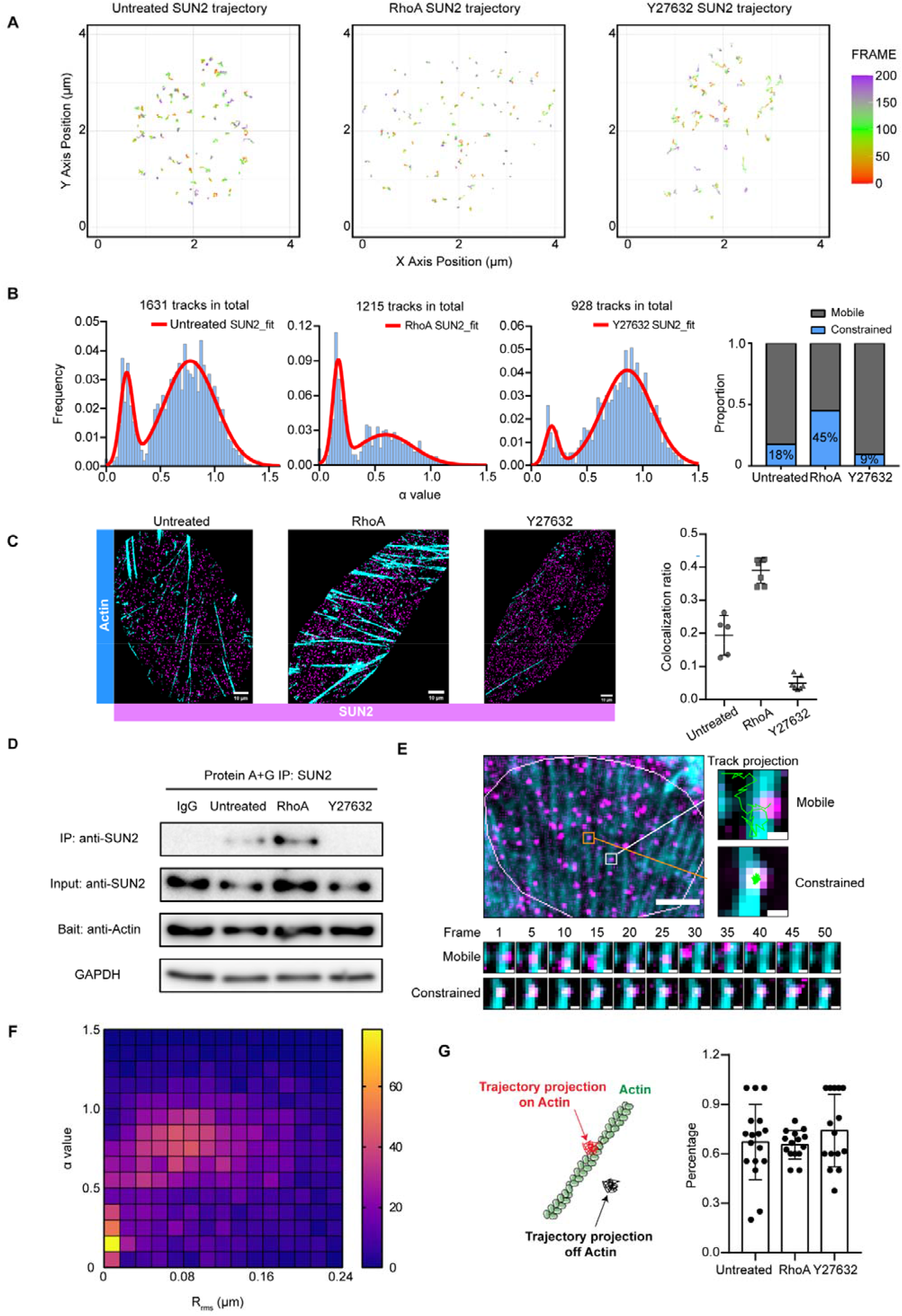
Actomyosin contractility regulates SUN2 dynamics. (**a**) Representative single-molecule tracking trajectories of SUN2 molecules in untreated, RhoA group, and Y27632 group. Scale bar, 1 μm. (**b**) α value histogram of SUN2 trajectories in untreated group (24 cells, 1631 tracks), RhoA group (19 cells, 1215 tracks), Y27632 group (18 cells, 928 tracks). Bin size, 0.025. Red line represents the fit curve. Right panel shows the proportion of constrained and mobile SUN2 subgroups. (**c**) 2D-SIM dual-color threshold images of actin and SUN2 molecules, Cyan: Fluorescence signal of actin. Megenta: Fluorescence signal of SUN2. Scale bar, 10 µm. Right panel shows the proportion of SUN2 molecules colocalized with actin bundles (median values, untreated, 0.22; RhoA, 0.413; Y27632, 0.04). (**d**) Immunoblots showing the immunoprecipitations between SUN2 and actin in untreated, RhoA group, Y27632 group. (**e**) Upper: Representative fluorescence images of single SUN2 molecules and actin bundles in live cell. White line represents the nucleus. Scale bar, 10□μm. Lower: Time lapse images of a mobile and a constrained SUN2 molecule. The exposure time was 10□ms and interval between each frame was 50□ms. Scale bar, 1□μm. (**f**) Heatmap of the average distance to the nearest actin bundle for each SUN2 trajectory and its corresponding alpha value. R_rms_ represents the Root Mean Square of the mean distance. Color bar represents the trajectory number. (**g**) (Left) Model of the location between constrained SUN2 trajectory projections and actin bundles. (Right) Quantification of actin colocalized SUN2 trajectory projections under different actomyosin contractility conditions.

We next aimed to probing the mechanism underlying the mechanical regulation of SUN2 dynamics. As SUN2 interacts with actin cytoskeleton through Nesprin-2G, we investigated whether this direct interaction contributes to the regulation of SUN2 dynamics under different mechanical conditions. Using Structure Illumination Microscopy (SIM), we measured the colocalization ratio between SUN2 and actin bundles near NE **(Fig. 2c and Extended Data Fig. 3d)**. Dual-color fluorescence labeling of actin and SUN2 in fixed cells showed that RhoA treatment induced formation of thick actin bundles near NE and Y27632 treatment disrupted those actin bundles in comparison with that in untreated cells **(Extended Data Fig. 3e-f)**. Accordingly, colocalization analysis indicates that the colocalization ratio between SUN2 and actin bundles was increased in the RhoA group and decreased in the Y27632 group **(Fig. 2c)**. It is worth noting that the colocalization ratio is very similar to the proportion of constrained subgroup in the corresponding single-molecule tracking data in **Fig. 2b**, implicating that the interaction between SUN2 molecules and actin cytoskeleton is key to regulate SUN2 dynamics on NE. In line with the imaging analysis, co-immunoprecipitation data also showed that the interaction between actin and SUN2 was positively correlated with the actomyosin contractility level **(Fig. 2d)**.

We performed dual-color single molecule imaging to provide direct evidence for the regulation of SUN2 dynamics by SUN2-actin interaction. By analyzing the trajectories of individual SUN2 molecules, we were able to obtain both of their mobility and spatial relationship with the Lifeact-EGFP-labeled actin bundles in living cells **(Fig. 2e and Supplementary Video 4)**. Interestingly, despite of the dramatic differed fractions of the mobile and constrained subgroups for different actomyosin contractility levels, most of constrained SUN2 molecules were found to be spatially correlated with the actin bundles with similar correlation levels (∼70%) under different actomyosin contractility conditions **(Fig. 2g)**. Correspondingly, when we calculated the root mean squared distance (R_rms_) for each individual SUN2 trajectory to its nearest actin bundle and correlated R_rms_ with the anomalous diffusion exponent α value, we clearly found that SUN2 molecules with small α values, *e*.*g*. <0.4, tend to colocalize with actin bundles (R_rms_ ≈0), and the α value is increased with R_rms_ **(Fig. 2f)**. These results strongly suggested that those less mobile SUN2 molecules are likely constrained *via* interaction with the actin bundles near the NE.

### 2.3 LaminA/C participates in regulating SUN2 dynamics

It was reported that SUN2 interacts with nuclear lamina with its N-terminus^25^. We wonder how its double-ended connections with nuclear lamina and actin cytoskeleton regulate the distribution and dynamics of SUN2. We first tracked SUN2 molecules in laminA/C and laminB1 KO cells, respectively **(Fig. 3a, Extended Data Fig. 4a and Supplementary Video 5-6)**. After MSD analysis, these data showed that compared to WT cells, cells lacking laminA/C had higher proportion of constrained SUN2 molecules, while laminB1 KO did not seem to affect SUN2’s dynamics **(Fig. 3c and Extended Data Fig. 4b)**. This observation is consistent with the previous report that SUN2 has stronger interaction with laminA/C than laminB1^20^.

**Fig. 3.**
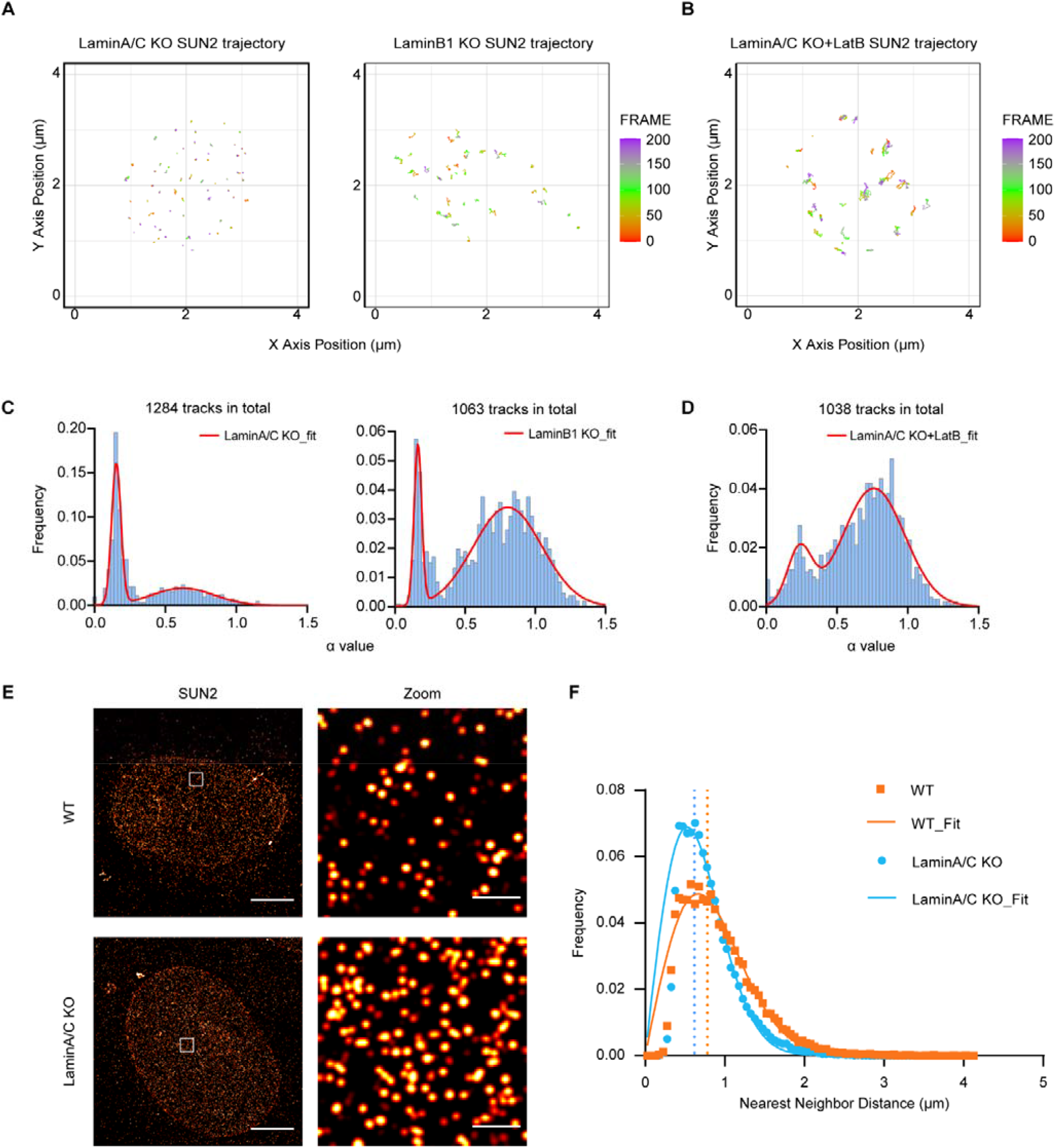
LaminA/C participates in regulating SUN2 dynamics. (**a, b**) Representative single-molecule tracking trajectories of SUN2 molecules in LaminA/C KO, LaminB1 KO cells (**a**) and LatB treated LaminA/C KO cells (**b**). Scale bar, 1 μm. (**c, d**) α value histograms of SUN2 molecules in LaminA/C KO group (16 cells, 1284 tracks) and LaminB1 KO group (14 cells, 1063 tracks) (**c**), and LatB treated LaminA/C KO cells (16 cells, 1038 tracks) (**d**). Bin size, 0.025. Red line represents the fit curve. (**e**) Representative STORM imaging of SUN2 under WT and LaminA/C KO conditions, respectively. Magnified views of the regions in the white boxes as right. Scale bar, 20 μm and 2 μm in the original and magnified images, respectively. (**f**) Histograms of the nearest neighbor distance for all clusters in WT and LaminA/C KO conditions. The dashed line represents the median value, WT: 0.7819 μm, LaminA/C KO: 0.6187 μm, bin size, 0.05.

Interestingly, we noticed that the α value distribution of SUN2 in laminA/C KO cells was similar with that in RhoA-treated cells **(Fig. 3c and 2b)**. STORM imaging also revealed similar SUN2 distribution in laminA/C KO cells and RhoA-treated cells **(Fig. 3e, f)**. Because laminA/C, unlike actin cytoskeleton, cannot actively exert force on SUN2, these data suggested that the SUN2-lamina connection passively counteracts with actomyosin contractility to regulate the distribution and dynamics of SUN2. Therefore, loss of laminA/C would make SUN2 be more constrained by actin cytoskeleton. In line with this hypothesis, when we added the actin polymerization inhibitor LatB to laminA/C KO cells to disrupt connections on both ends, SUN2 molecules became more dynamic **(Fig. 3b, d and Supplementary Video 7)**, and its α value distribution was very similar with that in Y27632 treated cells in which actomyosin contractility was lost while SUN2-lamina connection was preserved. Moreover, reinforced actomyosin contractility is known to unfold the Ig-domain of laminA/C into a soluble state^34^, which can free SUN2 from the lamina constraint. This may explain why laminA/C did not show resistance on SUN2 dynamics after strengthening actomyosin contractility. Taken together, these results suggested that actomyosin contractility is active and dominant in controlling SUN2 dynamics, while laminA/C is passive by docking SUN2 at the nuclear lamina.

### 2.4 Actomyosin contractility does not affect SUN2 oligomeric state

SUN2 was reported to present as monomeric and trimeric states on NE, and the matured trimer state is a structure prerequisite for its physiological functionating of SUN2^35^. In order to analyze whether actomyosin contractility affects the oligomeric state of SUN2, we applied the single molecule photobleaching assay to quantify the molecule number in individual SUN2 foci on NE **(Fig. 4a-f)**. The photobleaching steps indicated that most SUN2 foci (85-90%) were in trimeric state and there was no significant difference among the untreated, RhoA and Y27632 group **(Fig. 4g)**. Therefore, actomyosin contractility did not change the oligomeric state of SUN2.

**Fig. 4.**
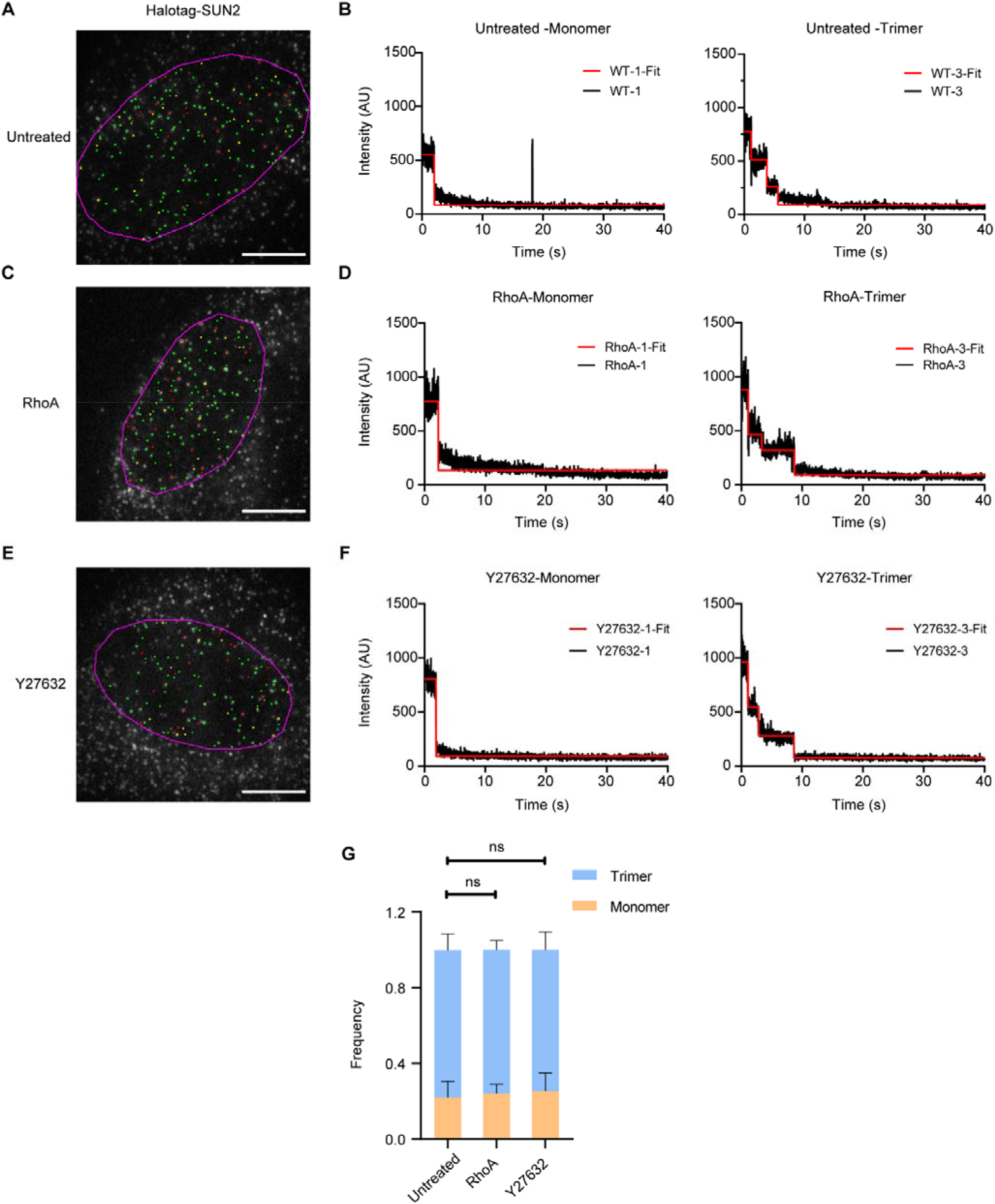
SUN2 oligomer states under different actomyosin contractility conditions. (**a, c, e**) Representative images of JF647-labeled Halotag-SUN2 under untreated (**a**), RhoA group (**c**) and Y27632 group (**e**). Magenta line indicates the nuclear periphery. SUN2 monomers are marked as red spots; SUN2 trimers are marked as green spots; Yellow spots represents the molecules without clear photobleaching steps due to frequent blinking. Scale bar, 20 μm. (**b, d, f**) Representative time traces of fluorescence intensity of SUN2 molecules in untreated (**b**), RhoA group (**d**), and Y27632 group (**f**). (**g**) Proportion of SUN2 oligomer states in untreated (22 cells, 3381 spots), RhoA group (18 cells, 1258 spots) and, Y27632 group (21 cells, 1944 spots).

### 2.5 Mechano-transduction through SUN2 contributes to perinuclear gene transcription

As a transducer for actomyosin contractility, we wonder whether SUN2 regulates the perinuclear gene transcription by transducing the external force. In order to dissect the mechanical response and decouple the subsequent biochemical effects, in addition to SUN2 KO, we designed two truncated mutants of SUN2 which are incapable of force transducing by detaching its connection to actin cytoskeleton *i*.*e*., ΔKashlid (569-585 aa) and to lamin, *i*.*e*., ΔLbd (Lamin binding domain, 1-139 aa), respectively. In line with the findings obtained by actin depolymerization using Y27632 in **Fig. 2a**, disconnecting SUN2 from actin cytoskeleton, *e*.*g*. the SUN2 mutant ΔKashlid, caused SUN2 molecules more mobile on NE than the WT group. Similarly, single molecule tracking of ΔLbd showed that the dynamics of this lamin-disconnected SUN2 mutant was more constrained **(Extended Data Fig. 5a)**, consistent with the observation in laminA/C KO cells in **Fig. 3c**. The single molecule tracking data of the WT group, RhoA group, Y27632 group, ΔKashlid, and ΔLbd together indicated that the physical linkage between actin cytoskeleton and lamina *via* SUN2 is required for force transduction.

To find out how SUN2 affects chromatin organization and gene transcription, we next investigated the distribution of several histone epigenetic modifications in SUN2 KO cells as well as cells separately expressing the two truncation mutants. While H3K4me3 and H3K27me3 were found to keep unchanged **(Extended Data Fig. 5b, c)**, the distribution of H3K9me3 changed clearly from a perinuclear enriched pattern in WT cells to an even distribution pattern in all three SUN2 deficient groups **(Fig. 5a, b and Extended Data Fig. 5d)**. These results strongly suggested that the integrity of the SUN2-containing LINC complex is essential for maintaining the perinuclear constitutive heterochromatin structure.

**Fig. 5.**
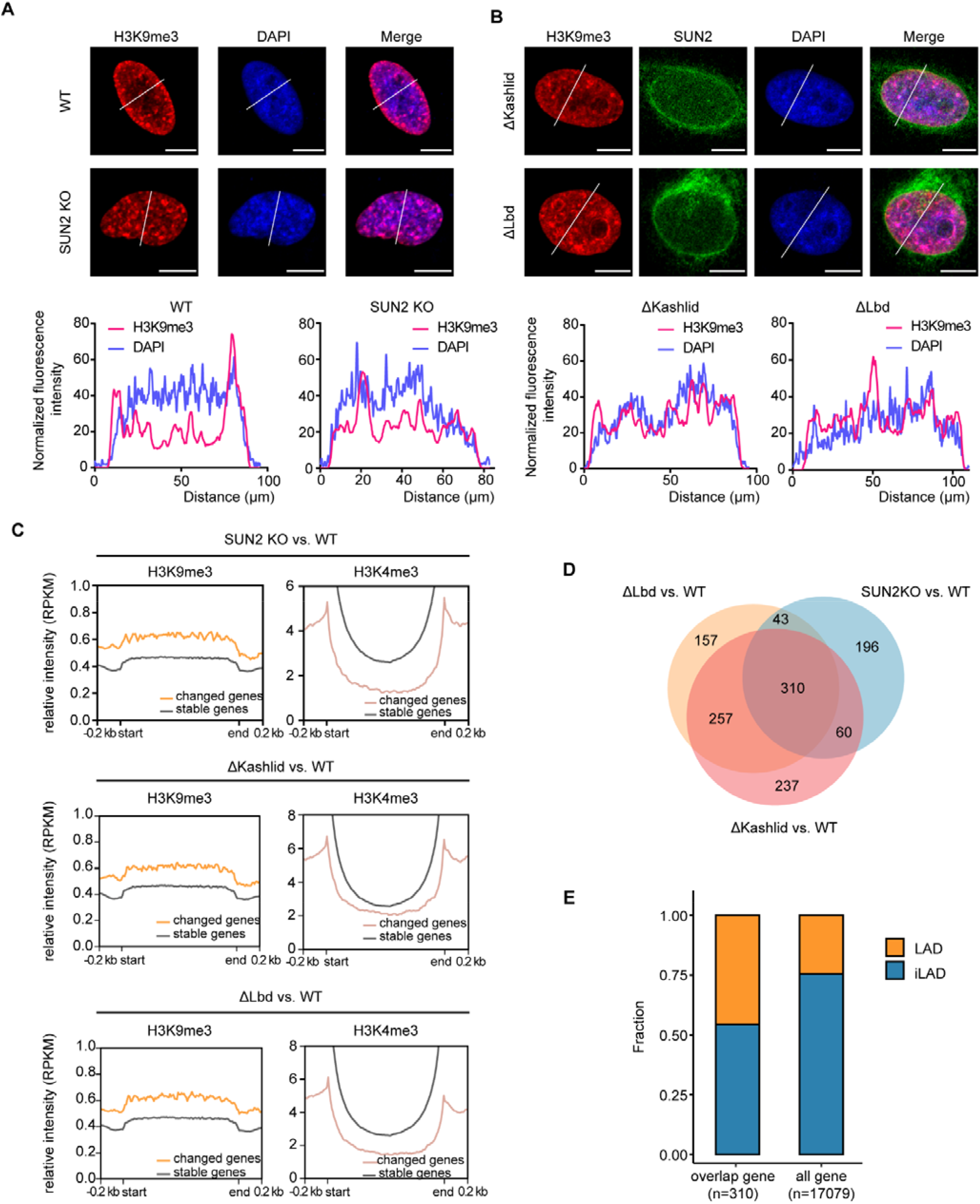
SUN2 participates in regulating perinuclear gene transcription. (**a**) Representative confocal images of H3K9me3 and DAPI in WT and SUN2 KO cells. Lower panel shows the intensity profiles marked by the white lines in the upper panel. Scale bar, 10 μm. (**b**) Representative confocal images of H3K9me3 and DAPI in ΔKashlid and ΔLbd cells. Lower panel shows the intensity profiles marked by the white lines in the upper panel. Scale bar, 10 μm. (**c**) Normalized reads intensity profiles of H3K9me3 and H3K4me3 signal around differentially expressed genes in SUN2 KO, ΔKashlid and ΔLbd groups. (**d**) The overlapped altered genes in SUN2 KO, ΔKashlid and ΔLbd cell groups. (**e**) The fraction of the altered genes in LADs and iLADs.

We then performed RNA-seq to investigate how the linkage between actin cytoskeleton and nuclear lamina through SUN2 regulates gene expression. In line with the imaging data, RNA-seq data showed that the genes with significant changes in expression levels between WT and SUN2 deficient cells were highly enriched in the H3K9me3 chromatin region, but rarely appeared within the H3K4me3 chromatin region **(Fig. 5c)**. Importantly, the genes that changed significantly are highly similar among the two truncation groups and SUN2 KO group. For instance, the ΔKashlid group and the ΔLbd group had ∼70% overlap of the changed genes, and both gene sets were half overlapped with the SUN2 KO gene set **(Fig. 5d)**. Given that the common character of SUN2 KO, ΔKashlid, and ΔLbd is the loss of physical linkage between actin cytoskeleton and nuclear lamina, it is tempting to speculate that the overlapped genes of the three groups are the genes that are directly regulated by mechanotransduction *via* the SUN2-containing complex as well as the genes that cells subsequently use to adapt to the mechanical changes. Indeed, GO analysis of the overlapped 310 genes showed most of the altered genes are involved in cell biomechanical processes such as cell migration and cell communication **(Extended Data Fig. 6a)**. Intriguingly, we found that the majority of the commonly altered genes were more enriched within the LAD regions **(Fig. 5e and Extended Data Fig. 6b)**. This finding is consistent with the facts that SUN2 interacts with the perinuclear lamina and SUN2 deficiency could induce perinuclear heterochromatin redistribution **(Fig. 5a, b)** and lamina ruffling **(Extended Data Fig. 6c)**. Because LADs and H3K9me3 regions are both typical cell-type invariant constitutive genomic regions near the nuclear envelope, these data indicated that SUN2 is a key element in maintaining the constitutive perinuclear chromatin, thereby regulating genes located in heterochromatin and LADs. These results implicate that proper connections with actin cytoskeleton and nuclear lamina are the basis for SUN2 to transmit mechanical signals and then regulate gene transcription.

## Discussion

LINC complex is a central structural component for nuclear mechanotransduction. Lack of high spatiotemporal studies have hampered the mechanistic understanding of LINC complex. Single molecule tracking and super-resolution imaging have become powerful methods for dissecting the distribution and dynamics of biomacromolecules, including those on the nuclear membrane, such as the nuclear pore complex^36^ and the lamin network^30,37^. In this study, we investigated the SUN2-containing LINC complex under different contractile forces at the single molecule level. We found that both the distribution and dynamics of SUN2 are dependent on the contractile force. Increased actomyosin contractility enriches SUN2 molecules from ER to NE and constrains their mobility. Further studies on drug treatments and truncated SUN2 mutants proved that the constraint on SUN2 required connections with both actin and laminA/C. We propose that lamin A/C and actin cytoskeleton form a perinuclear scaffold in which actin applies upward contractile force on the SUN2-containing LINC complex and lamin A/C passively counteracts the contractile force in a tug-of-war manner, and thereby co-regulates SUN2 distribution and dynamics on the nuclear membrane **(Fig. 6)**.

**Fig. 6.**
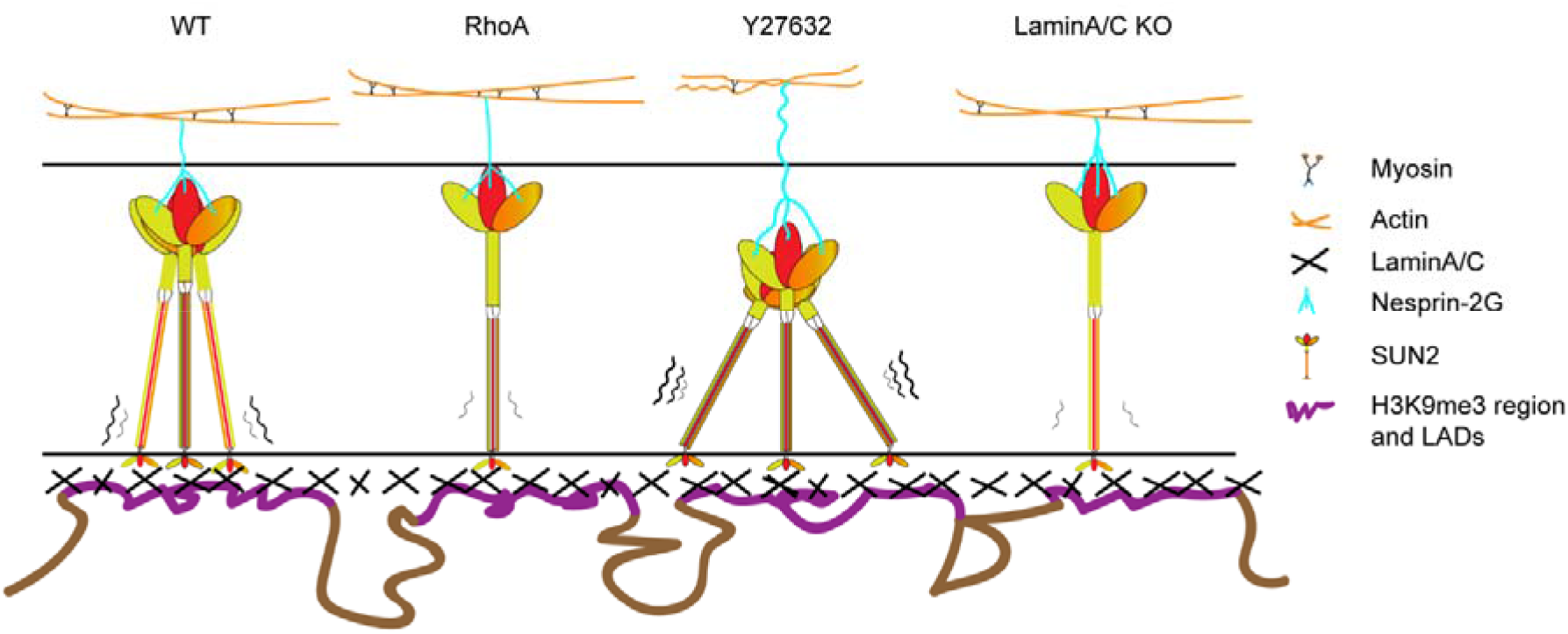
Model of actin and LaminA/C co-regulating SUN2 dynamics in a tug-of-war manner. Actin cytoskeleton and Lamin A/C form a perinuclear scaffold interconnected by SUN2. Actin applies upward contractile force which constraints SUN2 dynamics on the outer nuclear membrane. LaminA/C interacts with SUN2 and thereby passively counteracts the contractile force in a tug-of-war manner. Genes located in LADs and H3K9me3 enriched heterochromatin regions are more likely to be mechanically regulated through SUN2.

More specifically, our single molecule tracking data of SUN2 on the nuclear membrane revealed two mobility subgroups and the low mobility SUN2 molecules are clearly more associated with the actin than the high mobility ones **(Fig. 2)**. We also found that the actomyosin contractility can alter the proportion of the two subgroups. While several previous studies have observed the dynamic property of SUN2 by fluorescence photobleached recovery (FRAP) on the nuclear envelope^38,39^, they mainly described the diffusive capability of SUN2 between ER and nuclear membrane **(Fig. 1)**. Our single molecule tracking approach extends upon these prior findings by measuring the diffusion of individual SUN2 molecules which are further assigned to spatial correlation with actin cytoskeleton.

Actomyosin contractility has been found to perturbate the protein conformation of Nesprin-2G by loading an upward force^40^. It would be interesting to know whether and how SUN2 changes in protein conformation under actomyosin contractility. The crystal structure of SUN2 shows that the trans-luminal part of SUN2 is a coiled-coil domain that can be divided into coiled coil 1 (CC1) and coiled coil 2 (CC2), both of which consist of several folded subdomains^35^. The conformational changes of these hinge-containing subdomains make SUN2 fairly flexible in the NE luminal to buffer the force^41^. In addition, SUN2 exhibits as monomer or trimer on the NE referred to different maturation status. The trimeric state of SUN2 is known to be mainly regulated by several hydrophilic residues in the CC1 domain^42^. In this study, we found that the actomyosin contractility does not disturb the trimeric state of SUN2 on the NE by single molecule photobleaching experiments **(Fig. 4)**. Taken together, we speculate that actomyosin contractility cannot affect the trimerization of CC1 and CC2 may function as a buffering element to gauge the threshold that the SUN2-containing LINC complex transduces mechanical signals. Further studies with intramolecular FRET measurements would help to answer these questions.

Whether the force transmitted through the SUN2-containing LINC complex would affect gene transcription? While it was reported that Nesprin-2G depletion can inhibit biomechanical gene responses in murine myocardium, the direct mechanical response and subsequent biochemical effects were unclear^43^. In the current work, we designed two truncated SUN2 mutants to decouple the mechanical response from the subsequent biochemical effects. We observed that the genes with significant changes in expression are largely overlapped among SUN2 KO and the two truncated SUN2 mutants. Importantly, majority of the commonly altered genes are located in the LAD regions and involved in cell biomechanical processes, in line with the observations that SUN2 interacts with the perinuclear lamina and SUN2 deficiency can induce perinuclear heterochromatin redistribution. These observations collectively support that the SUN2-containing LINC complex plays it cellular functions mainly mechanically.

## Supporting information

Supplemental figures with corresponding titles and legends

## Author contributions

Y.S. and J.N. conceived and designed the research. J.N. performed the experiments. J.N. and X.W. analyzed the imaging results. W.Z. analyzed the RNA-seq results. Y.W. helps on the KO cell line generation. Y.Q. helps on the STORM imaging setup. X.H. helps on the SIM imaging. B.X. helps on the data processing discussion. C.L. helps on RNA-seq data processing discussion. Y.S. and J.N., wrote the manuscript.

## Acknowledgments

We thank Dr. Alan Jian Zhu’s and Dr. Xiaowei Chen’s lab for western blot exposure machine. We thank Dr. Luke Lavis for Halotag JF-650 dye. We thank the flow cytometry Core at National Center for Protein Sciences at Peking University, particularly Dr. Liying Du and Dr. Huan Yang, for technical help. We thank the imaging core facility at State Key Laboratory of Membrane Biology at Peking University for assistance with confocal microscopy, particularly Dr. Ye Liang, for technical help. This work is supported by grants from the National Science Foundation of China No.21825401 and the National Key R&D Program of China No.2022YFC3401100.

## Methods

### Cell lines and cell culture

Human embryonic kidney 293T (HEK 293T) cells were bought from ATCC (CRL-3216). Human breast cancer cell line MDA-MB-231 cells were bought from ATCC (HTB-26). Cells were cultured in Dulbecco’s modified Eagle medium (DMEM; Corning, 10-013-CRVC) with 10% fetal bovine serum (FBS; Gibco, 10099141C), 100 U/mL penicillin and 100 μg/mL streptomycin at 37 °C and 5% CO_2_ in a humidified incubator. Cells were passaged with the classical trypsin assay twice a week.

### Drug treatment

Rho activator II (Cytoskeleton, CN03) was used at the concentration of 1 μg/mL for 3-4 h before experiments. Y-27632 (Merck, Y0503) was used at the concentration of 10 μM for 1 h. Latrunculin B (Abcam, ab144291) was used at the concentration of 2.5 µM for 2-3 h before experiments. Actin-stain™ 488 Phalloidin (Cytoskeleton, PHDG1) was used at the ratio of 1:2000.

### Plasmids and transient infection

Full length of classical wild type human *Sun2* was cloned from the MDA-MB-231 cell-extracted cDNA library as the primary templet. Then EGFP-*Sun2* and Halotag-*Sun2* were subcloned into pcDNA3.1 and lentiviral vectors (lentiCas9-Blast) using Hieff Clone Plus Multi One Step Cloning Kit (Yeasen, 10912ES10). For SUN2 mutants, *ΔKashlid* (Δ569-585 aa) and *ΔLbd* (Δ1-139 aa) were amplified from the full length *Sun2* plasmids by circular PCR amplification, respectively. Lifeact sequence was self-made by annealing primers and cloned into pEGFP-N vector. All plasmids were transfected with Neofect (Neofect biotech, TF20121201) in accordance with the manufacturer’s protocol for 36–48□h.

### Generation of CRISPR/Cas9-mediated SUN2 and LaminA/C knockout cells

To generate knockout MDA-MB-231 cell lines, specific guide RNA sequences targeted to *Sun2* and *LMNA* (Designed by http://crispr.mit.edu) was ligated into the BbsI restriction site of the pX330-U6-Chimeric_BB-CBh-hSpCas9 (Addgene, 42230) vector and transfected in the wild type MDA-MB-231 cells, respectively. Cells were sorted by flow cytometry after 72 h transfection and finally validated by sequencing and immunoblot. LaminB1 knockout cell line was generated as described previously^44^.

### Generation of SUN2 related stable cell lines

To construct the SUN2 related stable cell lines without multiple isoforms interference, lentivirus plasmids lentiCas9-Blast (Addgene, 52962) containing: Halotag-*Sun2*, Halotag-*ΔKashlid*, Halotag-*ΔLbd*, EGFP-*Sun2*, EGFP-*Δ Kashlid* and EGFP-*ΔLbd* were each co-transfected with psPAX2 and pMD2.G in HEK293T cells. After 72 h, cell culture medium was collected and added to the SUN2 knockout MDA-MB-231 cells. After 72 h virus infection and selection by FACS, single cell clones were harvested for imaging a month later.

### Immunoprecipitation

Cells were lysed with lysis buffer (150□mM NaCl, 25□mM Tris-HCl, 0.5% NP-40, pH 7.4, 1□mM PMSF, 1□µg/ml leupeptin and 1□µg/ml aprotinin and protease inhibitor cocktail) for 30□min with gentle rotating, then centrifuged at 13000 g for 30 min at 4□°C for the supernatants. Protein A/G beads (Beyotime; P2012) were washed with lysis buffer and added to the primary Actin antibodies (10 ug/mL) followed by 1□h rotating at 4□°C. Mixture the antibody-incubated beads with supernatants (10 µL beads per 500 µL supernatants) by 4-5□h rotating at 4□°C and washed with lysis buffer three times. Finally, the beads were boiled in 1x SDS loading buffer, after 13000g centrifuge, the samples were separated by 8% SDS-PAGE gels and analyzed by western blotting.

### Western blotting

MDA-MB-231 cells were grown in 35□mm petri dishes. Cells were collected, washed twice with ice-cold PBS and lysed with 2 × SDS loading buffer, boiled for 5 min, and then centrifugation at 13,000 g for 10 min for the supernatants. Equal amounts of samples were loaded and then separated by SDS–PAGE on 8% gels and transferred onto a PVDF membrane by wet electrophoretic transfer. PVDF membrane was then blocked for 1 h in TBST (containing 5% skim milk). Primary and secondary antibodies were diluted in TBST (containing 2.5% skim milk). Primary antibodies were incubated overnight at 4°C overnight. After washed 5 min for three times with TBST, secondary antibodies were incubated at room temperature for 1 h. Images were acquired and analyzed by Odyssey imaging system (LI□COR biosciences) and ImageJ.

### Immunofluorescence staining

For all imaging assay, 35□mm glass-bottom microscope dishes (Cellvis, D35-20-1-N) were precoated with 5□μg/cm^2^ fibronectin (Sigma, F2006), and cells were seeded to achieve ∼60–80% confluency at the time of imaging. Cells were fixed with 4% PFA for 10 min, samples for STORM imaging were incubated by NaBH4 (1g/ml), permeabilized with 0.5% Triton in PBS for 5 min and then incubated in blocking buffer containing 5% BSA and 0.1% Triton for 30 min. Cells were then incubated with primary antibodies in blocking buffer overnight at 4□. After PBS washing for 3-5 times, then incubate with organic dyes-labeled secondary antibodies in blocking buffer for 1 h at room temperature. Finally, the labeled cells were washed again with PBS, then post-fixed with 4% PFA for 10 min. Actin in SIM images was stained by Actin-stain™ 488 phalloidin (Cytoskeleton, PHDG1) at 100 nM at room temperature for 30 min.

### Optical setup and image acquisition

Briefly, wide field images, STORM images and single molecule tracking images were taken on a custom Olympus IX83 inverted microscope equipped with a ×100 UPlanSApo, N.A.□=□1.40, oil-immersion phase objective, and Andor iXon Ultra EMCCD. The microscope stage incubation chamber was maintained at 37 °C and 5% CO_2_ for living cell. The laser power was modulated by an acousto-optic tunable filter (AOTF). We used highly inclined thin illumination (HILO) and carefully optimized the angle of the inclined light to reduce background from cell auto-fluorescence in the live cell tracking experiment. All confocal images were collected by a Zeiss 880 Airyscan (×100, N.A.□=□1.40). All SIM images were collected on a previously reported setup^45^.

For single molecule tracking, Halotag fused SUN2 variants cells were preincubated with 20 pM Halotag ligand JF650 for 10 min and washed with warm PBS for 3 times. After final wash, cells were replenished with phenol red-free medium for imaging. A 647-nm laser (∼10 μW) was used to excite JF650 with 10□ms exposure time for up to 2000 frames.

For STORM imaging, to get sparse labeling, cells were illuminated with 647□nm laser (∼300□mW) for 5–10□s to turn off most of the Cy5 labeled molecules. Then Cy5-labelled SUN2 were acquired up to 30,000 frames under excitation by a 647 laser at a power density of 3–5□kW□cm^−2^ and photoactivated with a 405□nm laser at a power density of 0.5□kW□cm^−2^ with 10□ms exposure time. Imaging buffer (100□mM Tris–HCl at pH□8.0, 20□mM NaCl and 10% glucose; all from Sigma-Aldrich) and an oxygen-scavenger system (60□mg ml^−1^ glucose oxidase, 6□mg□ml^−1^ β-ME (Gibco, 21985023)) were used for all STORM imaging experiments.

### Image analysis

All dynamic tracking image stacks was analyzed using Fiji’s Trackmate plugin (https://imagej.net/plugins/trackmate/) to obtain the trajectory of each single particle. The 2D time-lag mean squared displacement (TMSD) of each track was calculated by

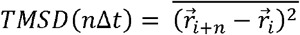

where ∆*t* is the time interval between frames, 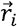 is the position of the particle at i-th frame in its trajectory, 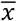 is the mean value of x. In practice, the trajectory is not necessarily continuous, that is, 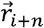 or 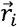may not exist, then this item will be directly skipped. The TMSD was fit to

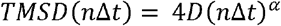

to calculate the diffusion coefficient (D) and anomalous diffusion exponent (α). In order to make the calculation results more accurate and reasonable, only the values of n from 0 to 5 in TMSD are used in the fitting.

For nearest neighbor distance (NND) analysis in STORM, the local maxima of gray value in the image is recorded as the preliminarily identified cluster position. Next, the images were binarized by Otsu algorithm. In order to prevent segmentation errors caused by extremely bright areas in the image, the gray value of the brightest 0.5% pixels in the image will be lowered before binarization. After binarization, foreground regions smaller than 4 pixels were considered as noise and discarded. According to the number of local maxima in each foreground region, all foreground regions were divided into two classes.

1. The number of local maxima in the foreground region is less than or equal to 1. The region is identified as a single cluster.
2. The number of local maxima in the foreground region is greater than 1. At this point, pixels in the region will be allocated to each nearby local maximum and form multiple clusters, and the allocation method of pixels and centroid position of each cluster will be optimized through iteration until the centroid positions movement between two iterations is less than the given threshold.

Then the centroid position of each cluster is calculated and the nearest neighbor distance between centroids is calculated. The statistical results were fitted by the following formula.

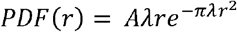

where A is the fitting parameter, which is related to the bin size of the histogram, r is the nearest neighbor distance, *λ* is the mean density of clusters.

For STORM image analysis, we used the Fiji’s plugin ThunderSTORM (https://github.com/zitmen/thunderstorm) for all drift corrections and image reconstruction under the authors’ protocol.

### Single molecule bleaching assay

Halotag fused SUN2 variants cells were preincubated with 20 pM Halotag ligand JF650 for 10 min. After 3 times wash, cells were fixed with 4% PFA. After fixation, cells were replenished with PBS for imaging. Cells were illuminated by a weak 647-nm laser with 10□ms exposure time until all the fluorophores are photobleached. Then the intensity changes of single molecules over time were recognized with 4×4 pixel square. Fitting to the bleaching steps was made by the home-written MATLAB codes.

### RNA-seq library construction

Total RNA was extracted from cultured MBA-MD-231 cells with 10,000,000 cells in each sample. RNA sequencing libraries were constructed using the NEBNext Ultra RNA Library Prep Kit for Illumina® (NEB England BioLabs). RNA-seq paired-end reads were sequenced on the Illumina NovaSeq 6000 platform.

### RNA-seq data analysis

Firstly, all the raw RNA sequences reads were cut adaptors and filtered using TrimGalore (https://www.bioinformatics.babraham.ac.uk/projects/trim_galore/) and mapped to human reference genome hg19 by STAR (v2.7.1a) with default parameters. High-quality mapped reads were quantified using htseq-count (v0.11.3) ^46^. Differentially expressed genes were analyzed by DEseq2 ^47^. The differentially expressed genes were defined by the cut-off: p valued < 0.05 and log2FoldChange ≥ 1 or log2FoldChange ≤ 1. Public ChIP-seq data used to do genomic distribution annotation of differentially expressed genes were download from ENCODE database (https://www.encodeproject.org/). The profile plots were prepared by deepTools (v3.5.0)^48^.

### REAGENT

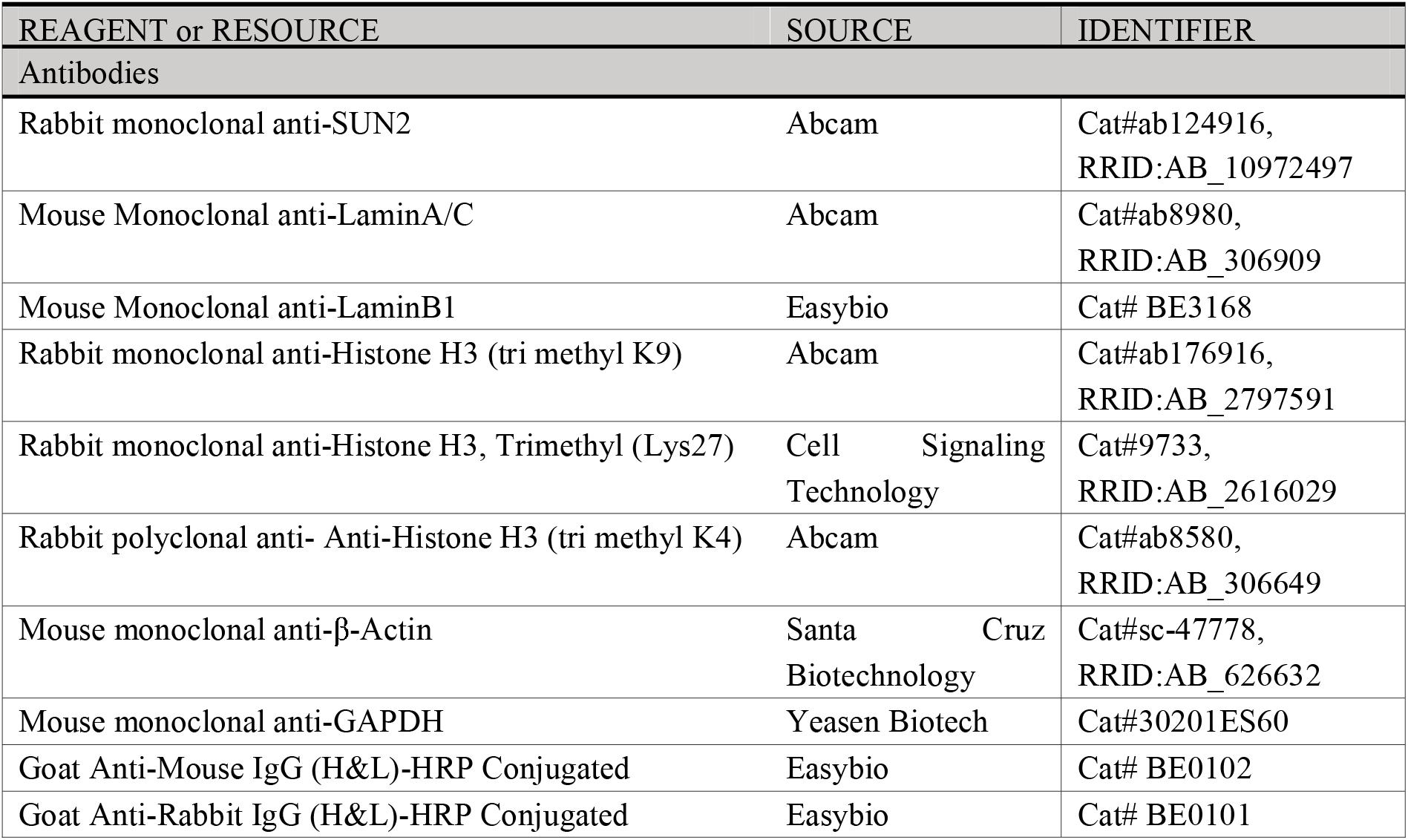
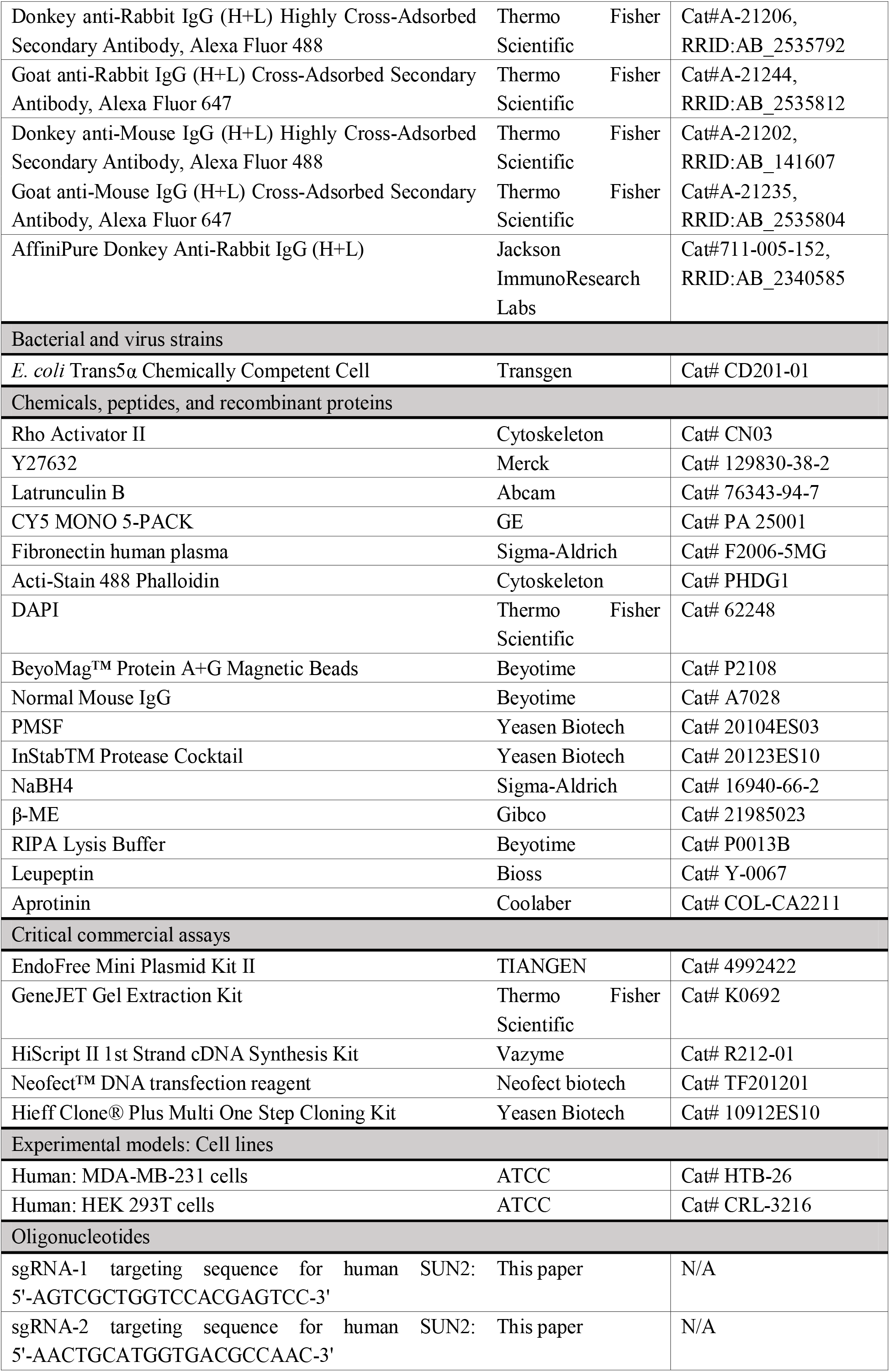
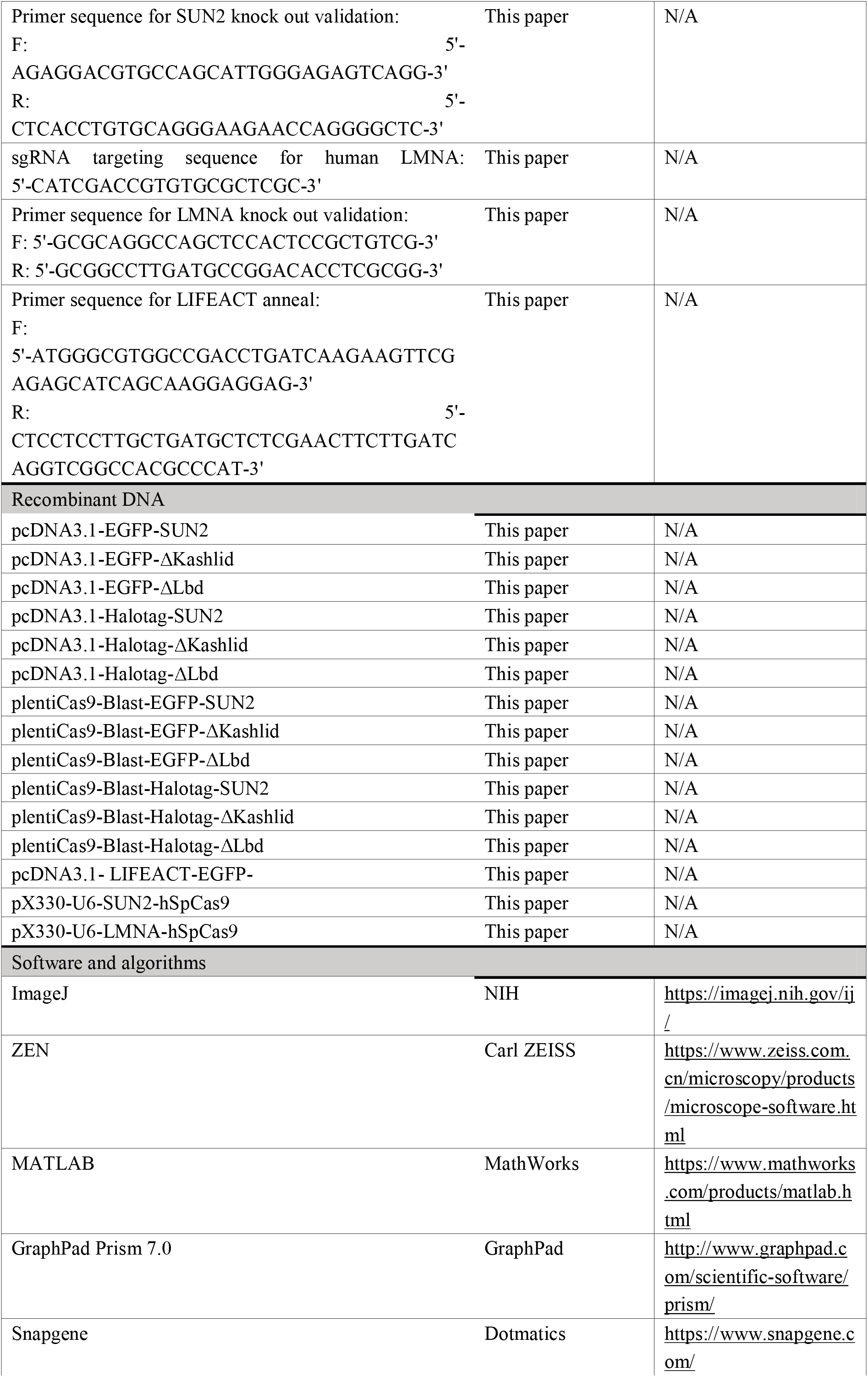
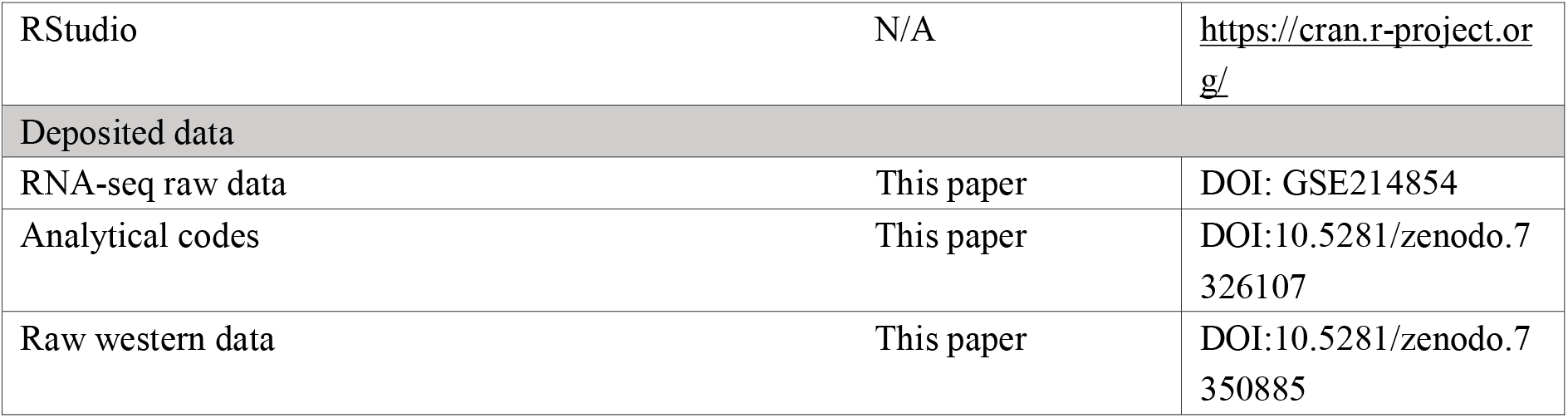

## Resource availability

### Lead contact

Further information and requests for resources and reagents should be directed to and will be fulfilled by the Lead Contact, Dr. Yujie Sun.

### Materials availability

All reagents generated for this paper are available upon request.

### Data and code availability

RNA-seq data have been deposited at GEO and are publicly available as of the date of publication. Accession numbers are listed in the key resources table. Microscopy data reported in this paper will be shared by the lead contact upon request. All original code and uncropped raw western images with molecular weight markers have been deposited at Zenodo and is publicly available as of the date of publication, DOIs are listed in the key resources table. Any additional information required to reanalyze the data reported in this paper is available from the lead contact upon request.

