## Supplemental figures with corresponding titles and legends for "Perinuclear force regulates SUN2 dynamics and distribution on the nuclear envelope for proper nuclear mechanotransduction"

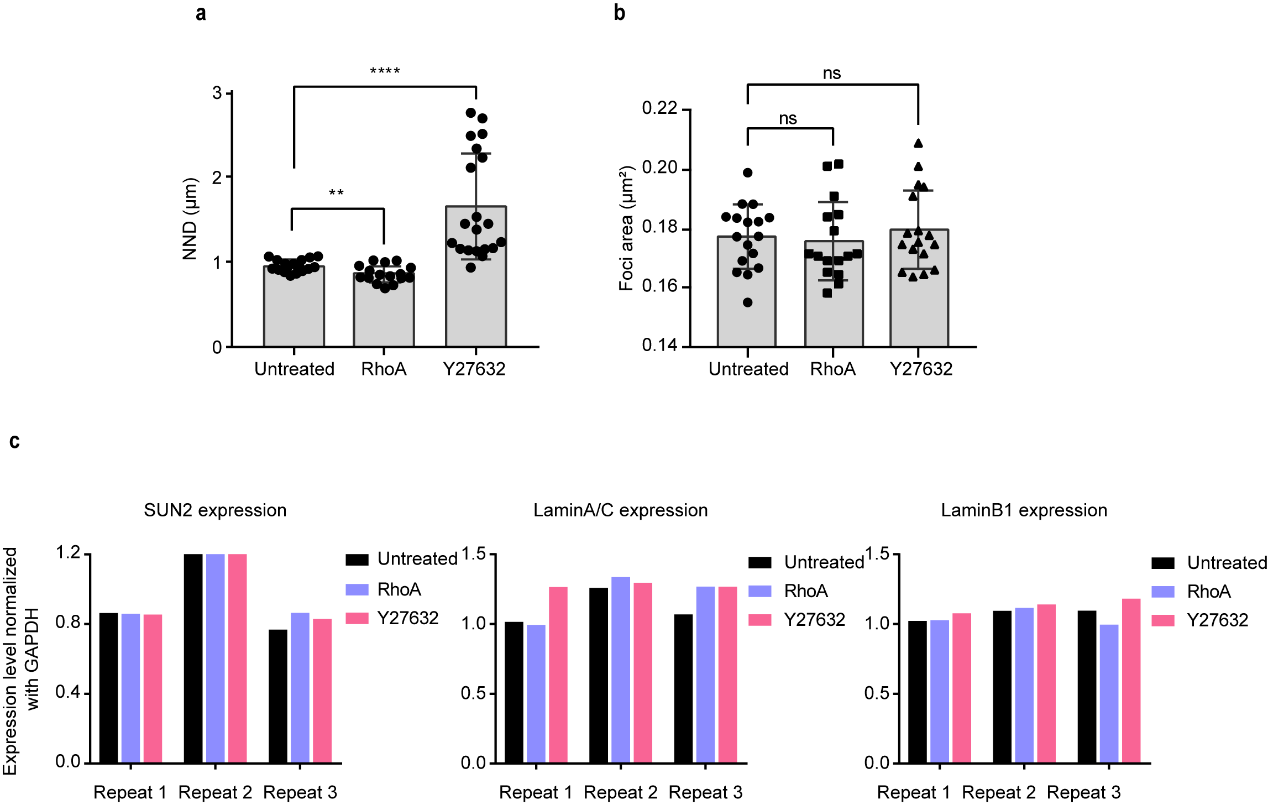

**Extended Data Fig. 1 NND and area of SUN2 foci in STORM images, Related to Fig. 1**

(**a**) Box-plot of NND of SUN2 clusters in each cell. Untreated (n=17 cells), RhoA (n=16 cells), Y27632 (n=20 cells).

(**b**) Box-plot of the area of SUN2 foci in each cell.

(**c**) Statistical results of whole cellular SUN2, LaminA/C and LaminB1 protein level under RhoA and Y27632 treatment for 3 repeats.
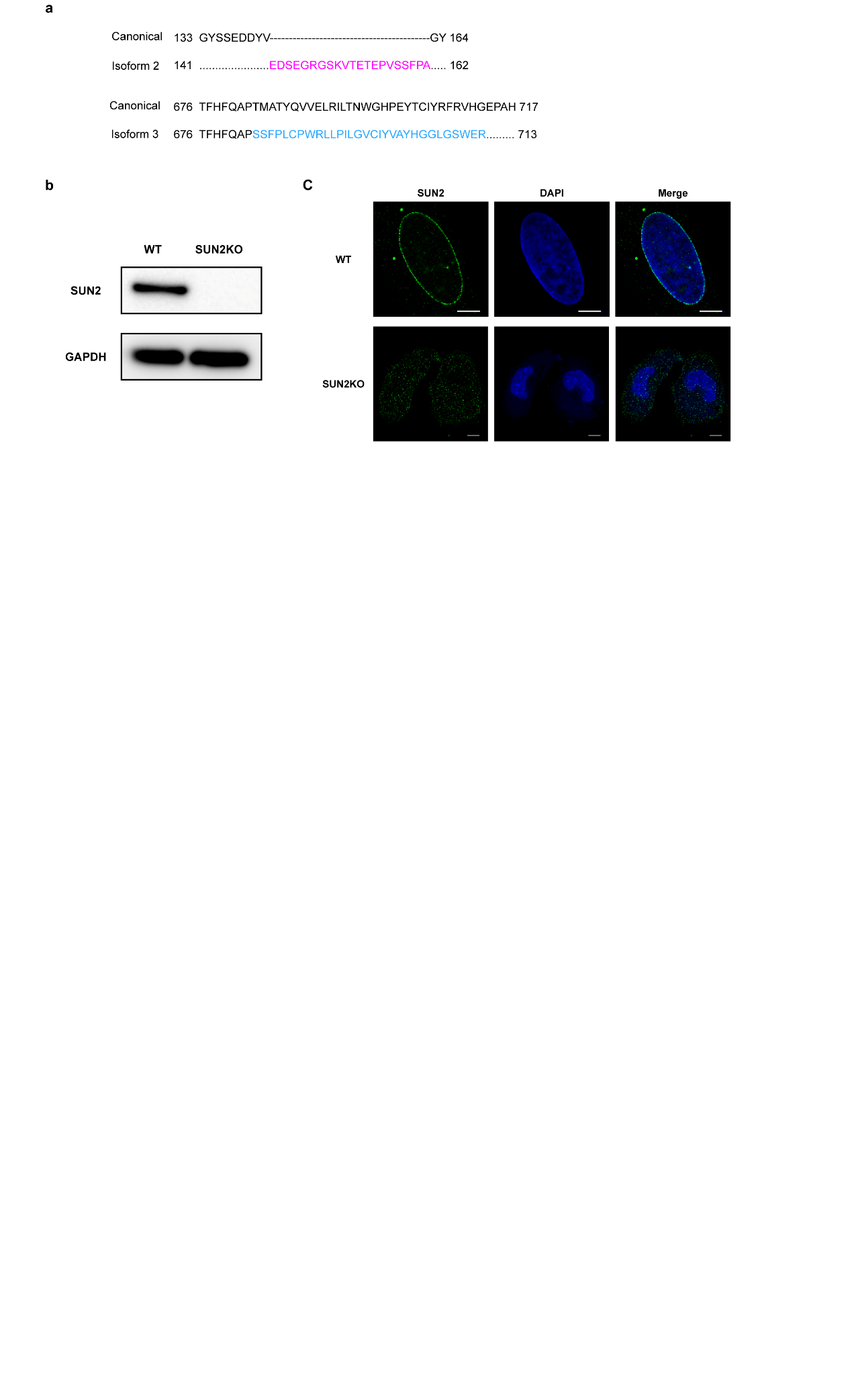

**Extended Data Fig. 2 Validation of SUN2 knockout, Related to Fig. 2**

(**a**) Typical Sun2 isoforms aligned with canonical Sun2 sequence.

(**b**) Immunoblot of SUN2 knockout MB-MDA-231 cells.

(**c**) Immunofluorescence of SUN2 knockout MB-MDA-231 cells. Scale bar, 10μm.

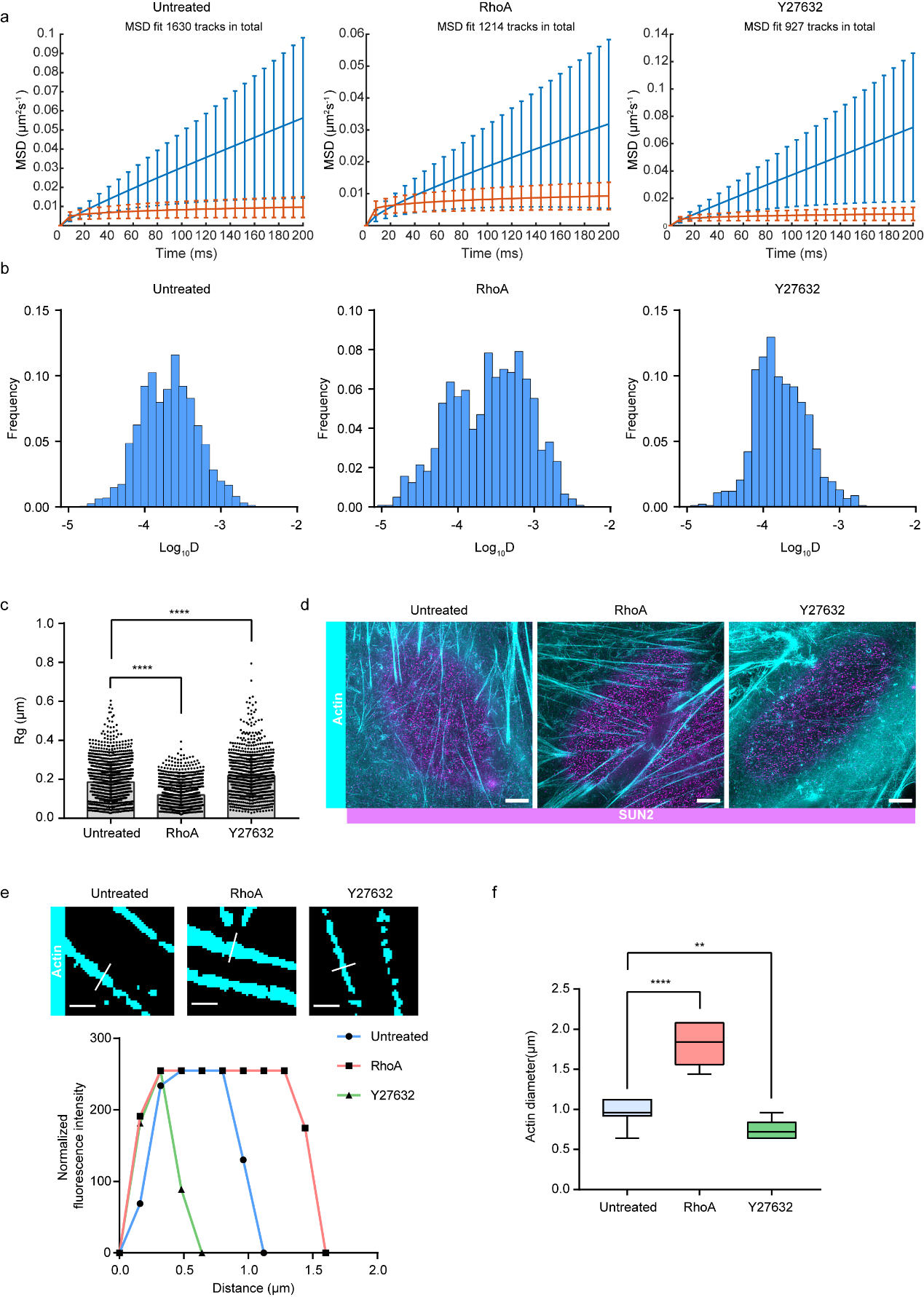

**Extended Data Fig. 3 Single molecule analysis of SUN2 molecules in three groups, Related to Fig. 2**

(**a**-**c**) MSD curve (**a**), diffusion coefficient (**b**) and radius of gyration (**c**) of untreated SUN2, RhoA group and, Y27632 group. ****, P < 0.0001.

(**d**) Colocalization of SUN2 and actin bundles under untreated, RhoA and Y27632 treatments by SIM imaging. Cyan: Fluorescence signal of Actin-488, Megenta: Fluorescence signal of SUN2-647. Scale bar, 10μm.

(**e**) Representative binary images of the actin bundle. Fluorescence intensity along the white line was measured using ImageJ software. Scale bar, 2μm.

(**f**) 10 random white line selections of actin diameter in each group. **, P < 0.01; ****, P < 0.0001.

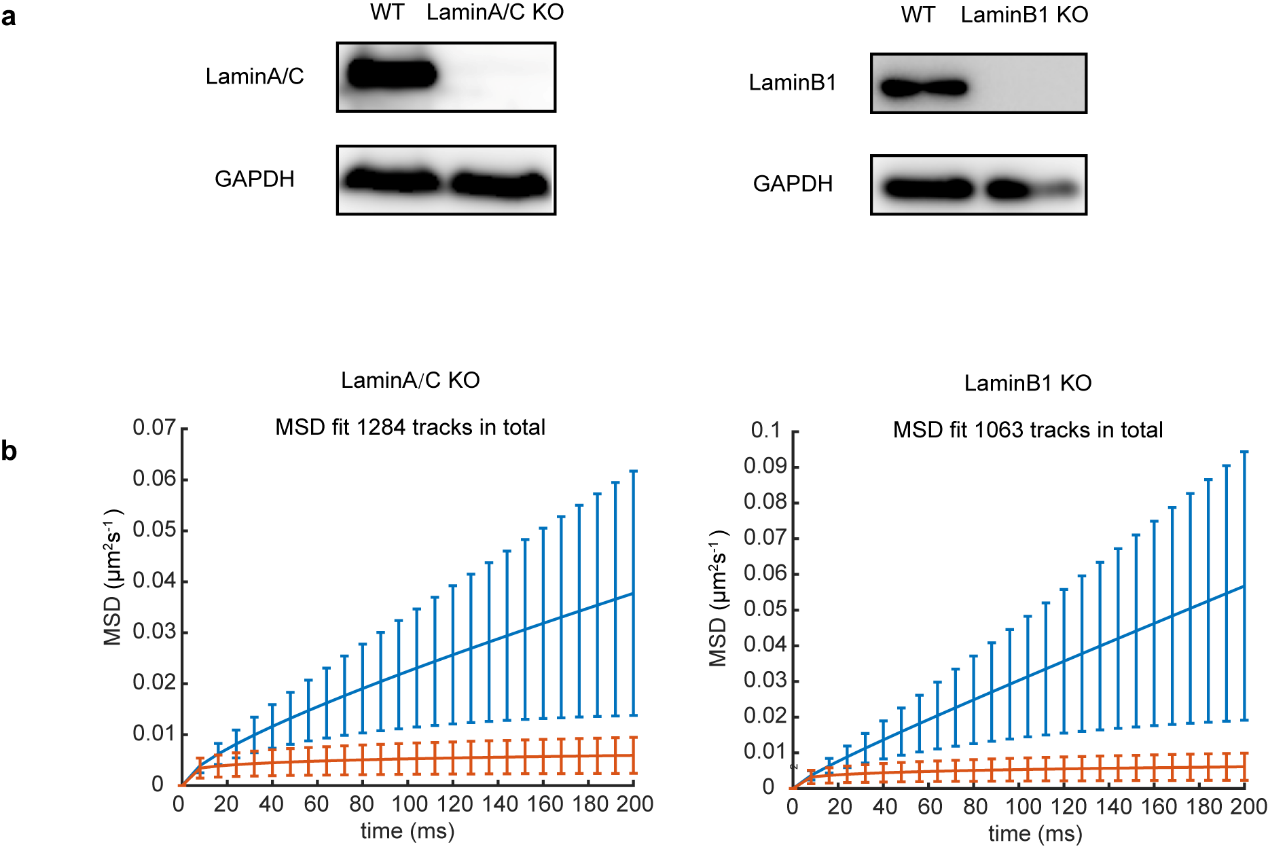

**Extended Data Fig. 4 MSD analysis of SUN2 molecules in LaminA/C or LaminB1 knockout cells, Related to Fig. 3**

(**a**) Immunoblot validation of LaminA/C and LaminB1 knockout.

(**b**) MSD curve of SUN2 molecules in LaminA/C or LaminB1 knockout cells.

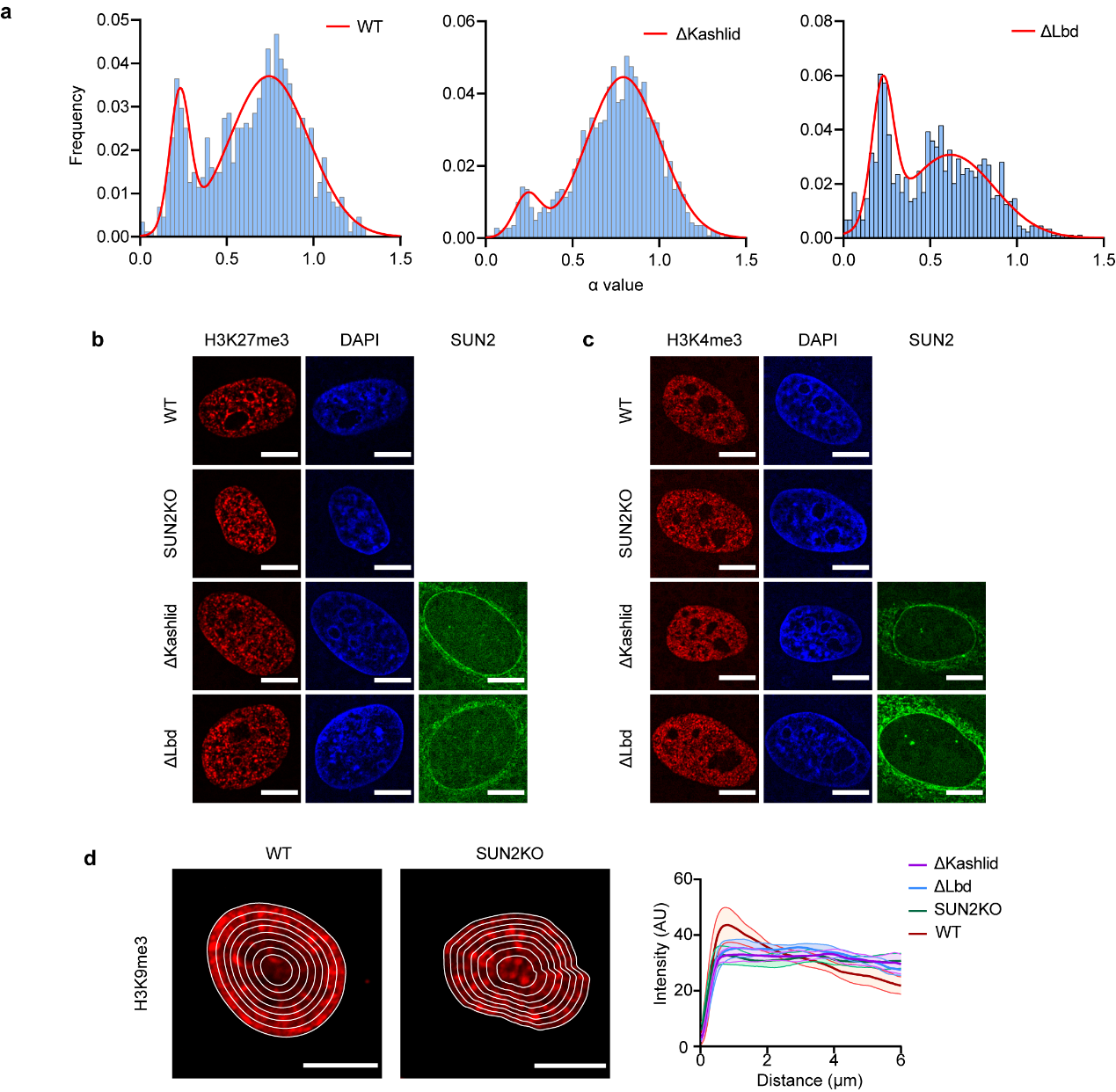

**Extended Data Fig. 5 Dynamic properties and chromatin regulation of SUN2 truncations, Related to Fig. 5**

(**a**) α value histogram of SUN2 trajectories in WT group (17 cells, 878 tracks), ΔKashlid group (20 cells, 1411 tracks), ΔLbd group (24 cells, 892 tracks), bin size, 0.025. Red line represents the fit curve. Right panel shows the proportion of constrained and motile SUN2 sub-group.

(**b**) Immunofluorescence of H3K27me3 in different SUN2 conditions. Scale bar, 10μm.

(**c**) Immunofluorescence of H3K4me3 in different SUN2 conditions. Scale bar, 10μm.

(**d**) Nucleus is divided into several shells with equal area form the nuclear periphery to interior. Representative averaged fluorescence intensity profiles of the concentric circles along the nuclear diameter in WT, SUN2 KO ΔKashlid and ΔLbd cells (n=10) stained with antibodies against H3K27me3. Scale bar, 10μm.

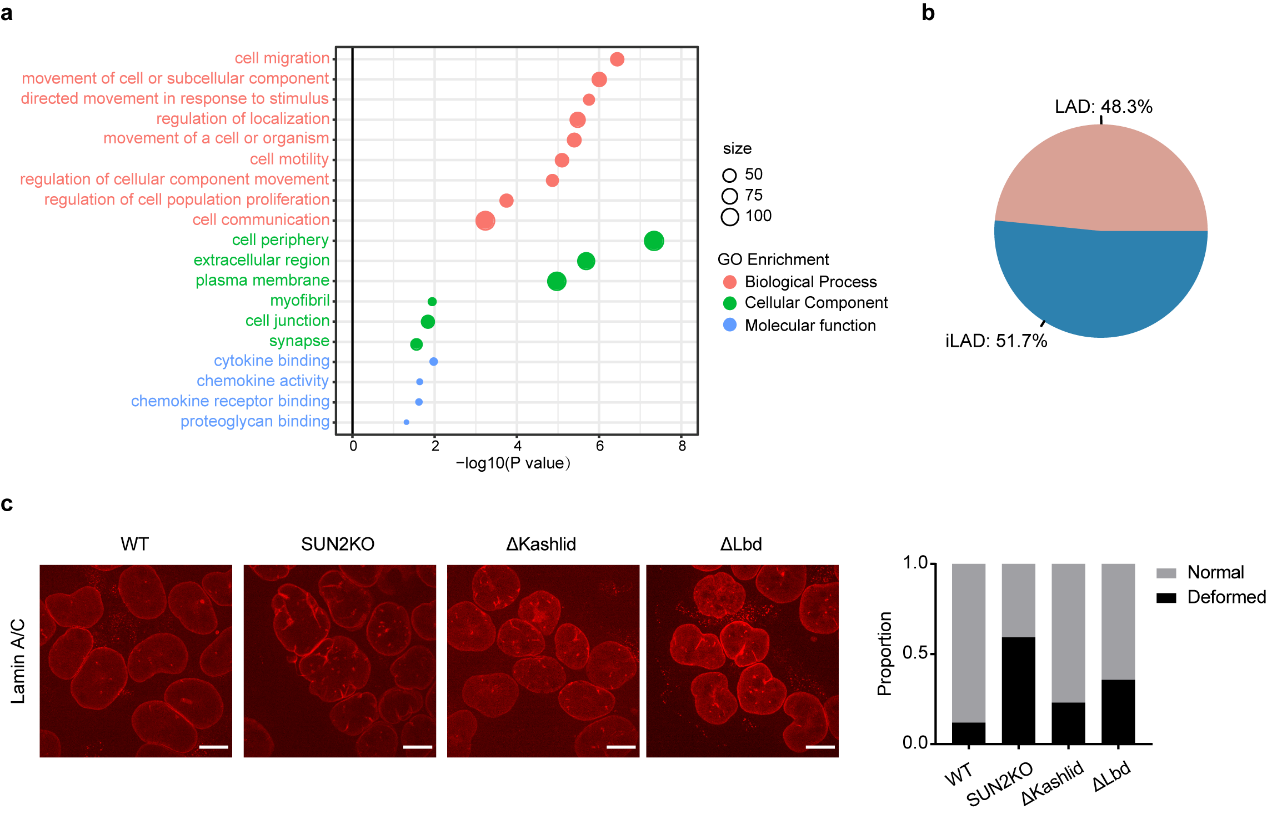

**Extended Data Fig. 6 GO analysis of SUN2 and NE wrinkles, Related to Fig. 5**

(**a**) GO analysis of the overlap genes.

(**b**) LAD types in MDA-MB-231 cells.

(**c**) Representative images of LaminA/C in WT, SUN2KO, ΔKashlid and ΔLbd cells. Box plot of the proportion of deformed and normal nuclear envelope in the cell population. Scale bar, 10μm.

**Supplementary Video legends**

**Supplementary Video 1. Single molecule tracking of untreated SUN2 molecules, related to Fig. 2**

Time-lapse movie of untreated Halotag fused SUN2 molecules. Exposure time: 10 ms, total frame: 200 frames. Scale bar: 10 μm.

**Supplementary Video 2. Single molecule tracking of RhoA treated SUN2 molecules, related to Fig. 2**

Time-lapse movie of RhoA treated Halotag fused SUN2 molecules. Exposure time: 10 ms, total frame: 200 frames. Scale bar: 10 μm.

**Supplementary Video 3. Single molecule tracking of Y27632 treated SUN2 molecules, related to Fig. 2**

Time-lapse movie of Y27632 treated Halotag fused SUN2 molecules. Exposure time: 10 ms, total frame: 200 frames. Scale bar: 10 μm.

**Supplementary Video 4. Dual-color live cell imaging of untreated SUN2 molecules and actin, related to Fig. 2**

Time-lapse movie of untreated Halotag fused SUN2 molecules (Megenta) and Lifeact labeled actin (Cyan). Exposure time: 10 ms, total frame: 200 frames. Scale bar: 10 μm.

**Supplementary Video 5. Single molecule tracking of SUN2 molecules in LaminA/C KO group, related to Fig. 3**

Time-lapse movie of Halotag fused SUN2 molecules in LaminA/C KO group. Exposure time: 10 ms, total frame: 200 frames. Scale bar: 10 μm.

**Supplementary Video 6. Single molecule tracking of SUN2 molecules in LaminB1 KO group, related to Fig. 3**

Time-lapse movie of Halotag fused SUN2 molecules in LaminB1 KO group. Exposure time: 10 ms, total frame: 200 frames. Scale bar: 10 μm.

**Supplementary Video 7. Single molecule tracking of LatB treated SUN2 molecules in LaminA/C KO group, related to Fig. 3**

Time-lapse movie of LatB treated Halotag fused SUN2 molecules in LaminA/C KO group. Exposure time: 10 ms, total frame: 200 frames. Scale bar: 10 μm.
